# Visuomotor Activation of Inhibition-Processing in Pediatric Obsessive Compulsive Disorder: A Magnetoencephalography Study

**DOI:** 10.1101/2020.11.23.394841

**Authors:** Eman Nishat, Colleen Dockstader, Anne L. Wheeler, Thomas Tan, John A. E. Anderson, Sandra Mendlowitz, Donald J. Mabbott, Paul D. Arnold, Stephanie H. Ameis

## Abstract

**Background:** The ability to inhibit a response is a component of executive control that is impaired in many individuals with obsessive compulsive disorder (OCD) and may contribute to clinical symptoms. This study explored whether neural processing during response inhibition, measured using magnetoencephalography (MEG), differed in a sample of medication-naïve youth with OCD, compared to typically developing controls (TDC).

**Methods:** Data was acquired from 20 medication-naïve children and youth with OCD (11.9 ± 2.1 SD years) and 14 TDC (12.3 ± 2.1 SD years). MEG was used to localize and characterize neural activity during a Go/No-Go task. Regional differences in amplitude of activity during a Go and No-Go condition between OCD versus TDC were examined.

**Results:** In response to the visual cue presented during the Go condition, participants with OCD showed significantly increased amplitude of activity in the primary motor (MI) cortex compared to TDC. A trend towards decreased amplitude of activity in the orbitofrontal cortex in OCD versus TDC was also found during stop errors in the No-Go condition.

**Conclusion:** Our preliminary study in a small medication-naïve sample suggests that neural response within motor and orbitofrontal regions may be altered during Go/No-Go Task performance, and that MEG as an imaging tool may be sensitive to detecting such differences.

## 1 Introduction

Obsessive compulsive disorder (OCD) is characterized by recurrent, intrusive thoughts and/or repetitive ritualistic behaviours (American Psychiatric Association, 2013). These symptoms are associated with significant distress, and deficits in occupational, academic, and social functioning (Wolff et al., 2017). The lifetime prevalence of the disorder is 2.3% in the general population and it affects up to 2% of children and youth (Ruscio et al., 2010; Fontenelle et al., 2006). Nearly half of adults diagnosed with OCD report the onset of symptoms prior to age 18 years (March et al., 2001). Earlier symptom onset has been associated with increased illness severity and persistence (Zijlmans et al., 2017). A focus on understanding brain processes linked to OCD in childhood and adolescence presents the opportunity to measure neural changes that may be directly associated with the disorder, rather than with potential confounds such as long-term medication exposure, illness duration, or learned strategies for behavioral compensation (Brem et al., 2012).

Impaired response-inhibition is a cognitive process that may contribute to the presence and maintenance of obsessive thoughts and compulsions (Yazdi-Ravandi et al., 2018). Response-inhibition is defined as top-down inhibitory control, limiting inappropriate response to external or internal stimuli (Berlin and Lee, 2018). A meta-analysis examining response inhibition performance across children and adults with OCD versus typically developing controls (TDC) demonstrated significant impairment in this domain in OCD (Hedge’s g = 0.77) (Lipszyc and Schachar, 2010). The frontal-striato-thalamic (FST) circuit, comprised of the supplementary motor area (SMA), orbitofrontal cortex (OFC), projections from the anterior cingulate cortex (ACC), white matter tracts including the anterior corpus callosum, cingulum bundle, and anterior limb of the internal capsule and subcortical regions, such as the striatum, subthalamic nucleus, and thalamus are implicated in habitual control and response inhibition (Morein-Zamir and Robbins, 2015). Among FST circuit regions, the ACC and OFC may be particularly important for response inhibition and relevant in OCD based on the role of the ACC in supporting cognitive control and error-monitoring, and the role of the OFC in emotional control, such as selective judgement or weighing of consequences (Fitzgerald et al., 2011; Abe et al., 2015).

A meta-analysis of task-related functional magnetic resonance imaging (fMRI) studies (n = 2345, mean age = 31.9 years) suggested that response inhibition elicits altered task-evoked brain response in adults with OCD compared to controls, including greater ACC activation (Rasgon et al., 2017). A limited number of neuroimaging studies have examined links between brain structure or function and response inhibition in pediatric samples (Norman et al., 2016). Available fMRI studies exploring response inhibition and related cognitive tasks have found both increased and reduced activation in OFC, medial prefrontal cortex, ACC, motor regions and caudate nucleus during task performance in children and youth with OCD versus TDC (Rubia et al., 2010; Fitzgerald et al., 2018; Fitzgerald et al., 2010; Britton et al., 2010). Studies have thus far included both medicated and un-medicated children and youth with OCD. However, medication exposure may be an important confound for consideration given recent findings that selective serotonin reuptake inhibitors (SSRIs), commonly prescribed for treatment of OCD may have effects on brain structure, function, and neurochemistry (Bernstein et al., 2019; Boedhoe et al., 2017). SSRIs have also been associated with brain change in children undertaking medication treatment targeting anxiety symptoms (Gaffrey et al, 2013).

Most neuroimaging studies in children and youth with OCD have focused on structural differences or have characterized neural activity in OCD using fMRI. However, magnetoencephalography (MEG) may offer some advantages over fMRI as it records neuromagnetic activity and enables tracking of neural activation with high spatial and temporal resolution (Ahlfors and Mody, 2019). Additionally, MEG can characterize the frequencies in which large groups of neurons fire during a particular task, including oscillations in the: delta (0.5-3 Hz), theta (4-7 Hz), alpha (8-12 Hz), beta (13-29 Hz), and gamma (30+ Hz) bandwidths (Todd et al., 2014). Thus far, very few studies have used MEG to study neural response in OCD. We are aware of just three MEG studies in children and youth (Amo et al., 2006; Korostenskaja et al., 2013; Mogadam et al., 2019). One of these available MEG studies in children and youth examined neural response to a cognitive task and found increased amplitude of prefrontal cortex response during a cognitive flexibility task in participants with a primary clinical diagnosis of OCD versus those with autism spectrum disorder or attention deficit hyperactivity disorder (n = 88, ages 8 to 15 years) (Mogadam et al., 2019). We are not aware of any MEG study in children and youth that has examined neural response in a medication-naïve sample.

The current study is the first, to our knowledge, to use MEG to explore differences in neural activity during response-inhibition task performance in a sample of medication-naïve children and youth with OCD compared to TDC. It was hypothesized that children and youth with OCD would feature increased amplitude of neural response within FST regions during response inhibition performance compared to TDC.

## 2 Materials and methods

### 2.1 Participants

All participants were recruited from The Hospital for Sick Children in Toronto. Informed consent was obtained from all parents and assent from all participants in accordance with the Declaration of Helsinki, and the study was approved by the Hospital Research Ethics Board. Structural MRI, MEG at rest and during Go/No Go Task performance, and behavioral data were acquired in 20 medication-naive children and youth (ages 8 to 16 years, mean age = 11.9 years ± 2.1 SD, 11M/9F), diagnosed with OCD by a child psychiatrist (PDA, SHA) or clinical psychologist (SM) in accordance with the Diagnostic and Statistical Manual of Mental Disorders (DSM) criteria, and 14 TDC (ages 8 to 16 years, mean age = 12.3 years ± 2.1 SD, 7M/7F). All participants were righthanded. Exclusion criteria consisted of prior psychopharmacological treatment exposure, a history of chronic neurological disorders, any previous serious head injury resulting in loss of consciousness, history of bipolar disorder, psychosis, or schizophrenia spectrum disorder in participants with OCD, or any history of psychiatric disorder based on parent report in TDC.

### 2.2 Clinical characterization in OCD

The diagnosis of OCD was confirmed using the Schedule for Affective Disorders and Schizophrenia for School-Age Children-Present and Lifetime version (K-SADS-PL) (Kaufman et al., 1997). The severity of obsessive-compulsive symptoms in children and youth with OCD was assessed using the Children’s Yale-Brown Obsessive-Compulsive Scale (CY-BOCS) and the Toronto Obsessive-Compulsive Scale (TOCS) (Park et al., 2016). The CY-BOCS is a 19-item, clinician-rated instrument with items 1-5 assessing the severity of obsession symptoms and items 6-10 assessing the severity of compulsion symptoms, the sum of which make up the total score (total score <13 = mild, 13-22 = moderate, >22 = severe OCD symptoms) (Scahill et al., 1997). The TOCS is a 21-item parent or self-report questionnaire, adapted from the CY-BOCS, providing additional quantitative information on obsessive-compulsive traits in children and youth (Park et al., 2016).

### 2.3 Magnetic resonance imaging

MRI scans were obtained for all participants on a 3.0T Siemens TIM Trio scanner, T1-weighted images were acquired using a 3D MPRAGE Grappa 2 protocol (TR/TE = 2300 per 2.96 ms, voxel size 1.0 mm x 1.0 mm x 1.0 mm). During the scan, children were positioned to watch videos via goggles to minimize head motion. Fiducials placed prior to MEG scanning were kept in place during the MRI for later co-registration.

### 2.4 Magnetoencephalography recordings

A whole-head 151 channel CTF MEG system placed in a magnetically shielded room was used to record neuromagnetic activity. MEG data were collected continuously at a rate of 1200 samples per second and were filtered at 0.3 to 300 Hz. Before data acquisition, each participant was fitted with one fiducial localization coil placed at the nasion and two placed at the preauricular points to localize the position of the head relative to the MEG sensors. Participants lay supine with their eyes open and fixated on a black cross (2 x 2 cm) on a semi-transparent screen placed 50 cm from their eyes. To monitor eyeblinks and saccades, electrooculograms were placed distal to the lateral canthus of each eye, and one on the left mastoid process. To monitor head movement, a headtracking system was used during data acquisition. The time points of overt saccades or eye blinks were visually inspected and trials including eye blinks prior to cue onset were excluded.

### 2.5 Go/No-Go task

The Go/No-Go task is a cognitive task assessing the ability to withhold (inhibit) a response when presented with a No-Go signal (i.e, signal to withhold response). The Go condition is a low cognitive load task that probes visuomotor engagement when a Go cue (green cross) is presented. In the No-Go condition, both a Go cue (green cross, cue to respond quickly) and No-Go cue (red cross, cue to withhold response) is presented (Georgiou and Essau, 2011). The No-Go condition is considered to have a higher cognitive load than the Go condition due to the need to discriminate between stimuli and cognitively select for an appropriate response (Dockstader et al., 2014).

### 2.5.1 Go task

As described in detail by Dockstader et al. (2014), during the Go task (serving as a control condition), participant’s eyes were open and fixated on a black cross. Their dominant hand was resting at their side with their thumb resting on a button box response button. The black cross was pseudo-randomly replaced with a green cross, temporally jittered between 1.5-2.5s and accompanied by a static visual contrast grating in the lower visual field. The location and dimensions of this contrast grating have been shown to elicit a strong visual evoked field approximately 75ms after cue onset (Koelewijn et al., 2011). During Go trials, participants were instructed to press the button with their dominant thumb immediately following the presentation of the green cross. The timing of the button-press in response to the green cross was recorded (Go reaction time). Each participant underwent 100 Go task trials (Dockstader et al., 2014).

### 2.5.2 No-Go task

The No-Go task served as the inhibitory condition. During the No-Go task, the black fixation cross was replaced with either a green or red cross accompanied by a static visual contrast grating. Participants were instructed to press the button immediately following the presentation of the green cross and were instructed to inhibit a button-press following the presentation of the red cross. Reaction time was recorded as the timing of button-press in response to the green cross. Stop errors were recorded as any button-press following a red cross (i.e., No-Go signal). There were 197 green crosses and 100 red crosses presented, pseudo-randomly, such that 67% of trials were green crosses (Go cues) and 33% were red crosses (No-Go cues). The ratio of Go to No-Go stimuli was selected so that the majority of trials were Go trials (requiring a button-press). This required participants to inhibit the tendency to respond during the No-Go trials, in keeping with previous use of the Go/No-Go Task in a pediatric sample (Dockstader et al., 2014; Vara et al., 2014; Vidal et al., 2012). The presentation order was counterbalanced across all participants in both groups.

### 2.6 Brainstorm data pre-processing

MEG datasets for all participants were processed and analysed with Brainstorm (Tadel et al., 2011), which is freely available for download under the GNU general public licence (http://neuroimage.usc.edu/brainstorm). Data were down sampled to 600 samples per second prior to processing. For both Go and No-Go conditions, datasets corresponding to the timing of visual cue were defined with an epoch time of −400 ms to 1000 ms, as a visual response is expected to occur within this time range. DC offset, which is signal noise that occurs at cue onset, was removed and a baseline epoch was defined as −100 ms to 0 ms. The datasets corresponding to time of expected button-press response (i.e., response to Go cue and stop errors to No-Go cue) were defined with an epoch time of −300 ms to 300 ms, around the motor response. This parameter allowed for analysis of any activity occurring prior to or following the button-press. DC offset was removed using the baseline computed for each output file and bandpass filtered from 1-40 Hz, with a Notch filter of 60 Hz (Tadel et al., 2011). MEG data was co-registered with MRI data to create high resolution, three-dimensional, differential images of activations for Go and No-Go trials, relative to baseline.

### 2.7 MEG analysis

#### 2.7.1 Source localization and extraction of response amplitude data

Latency of visual response in the average group waveform activity across sensors in both OCD and TDC groups in Go and No-Go conditions was first identified. The time of peak activity, at ~100 ms (expected time of visual response) was recorded as the latency of visual response to the visual cue in each group. Maximum amplitude of activation at peak timepoints in the group average waveforms for each group (OCD and TDC) for Go button-press and No-Go stop errors were also identified. An average activation map for each group (OCD and TDC) was created for Go buttonpress and No-Go stop errors by subtracting the baseline epoch from the active epoch, with a linear minimum norm estimation algorithm applied (Tadel et al., 2011). Average activation maps for each group (OCD and TDC) were used to visualize and identify any regions of activation corresponding with peak time points (derived from the group waveform) for both Go button-press and No-Go stop errors. Subsequently, response amplitudes and standard deviation were extracted at the participant level based on peak activation between 100 ms to 300 ms, as peak activation for group waveforms was within this time window (for both Go button-press and No-Go stop errors). We confirmed regions of activations for each individual were the same as the regions activated at the average group level by creating activation maps corresponding with peak time points for individual participants.

#### 2.7.2 Statistical analyses

Two-sample *t*-tests were used to examine between-group differences in Go (amplitude of activation and reaction time during button-press) and No-Go task performance (amplitude of activation during stop errors and number of stop errors). Effect sizes (Cohen’s d) were calculated to evaluate the magnitude of any significant between-group differences. Given the small sample explored in this preliminary MEG analysis, p < 0.05 was considered significant andp-values were not adjusted for multiple comparisons.

The Multivariate Imputation by Chained Equations (MICE) package on R was used to interpolate random missing values due to missing markers indicating time of cue presentation in dataset files or missing clinical scores (R Core Team, 2018). Statistical outliers in amplitude of activation were removed using the Box Plot Statistics function in the R Graphics Devices (grDevices) package from calculation of correlation analyses (R Core Team, 2018).

#### 2.7.3 Exploratory post-hoc analysis

Time-frequency response. In order to explore the frequencies driving the responses during the Go and No-Go task conditions, time-frequency response (TFR) plots were generated for regions of interest (i.e., within the FST circuit) that differed between groups during source localization. A Morlet wave analysis of changes in oscillatory frequency was used by selecting for virtual sensors associated with regions of interest. A reduction in frequency oscillations was specified as event-related desynchrony (ERD), whereas an increase in frequency oscillations was specified as event-related synchrony (ERS). Peak frequency oscillations were identified in delta (0.5-3 Hz), theta (4-7 Hz), alpha (8-12 Hz), beta (13-29 Hz), and gamma (30+ Hz) frequency bands. Time-frequency activity was analyzed relative to baseline activity (activity prior to cue onset at 0 ms).

Correlation of event-related amplitude with clinical scores. Exploratory correlation analysis was conducted to examine whether FST regions of interest that differed between groups during response inhibition performance were associated with clinical symptoms. Correlations were computed using Pearson correlation on R Statistical Analysis (R Core Team, 2018).

## 3 Results

### 3.1 Participants

The demographic characteristics of the total sample by group are presented in Table 1. Based on the CY-BOCS, average symptom severity in participants with OCD was in the high moderate range (22.84 ± 4.5 SD). Data was adequate across participants with respect to head motion and apparent perception of cue (i.e., no participant scan included head movement ≥ 5 mm, or overt saccades or eye movements occurring between −200 to 0 ms prior to cue onset). Two OCD participants were removed due to missing MRI scans, and one was removed due to missing buttonpress markers in MEG data files and no available datapoints to perform imputation. The total number of participants analyzed in the OCD group was n = 17 in the Go condition and n = 14 in the No-Go condition as three participants did not make any errors. The total number of participants analyzed in the TDC group for both Go and No-Go conditions was n = 13, due to missing data in one participant.

**Table 1.**
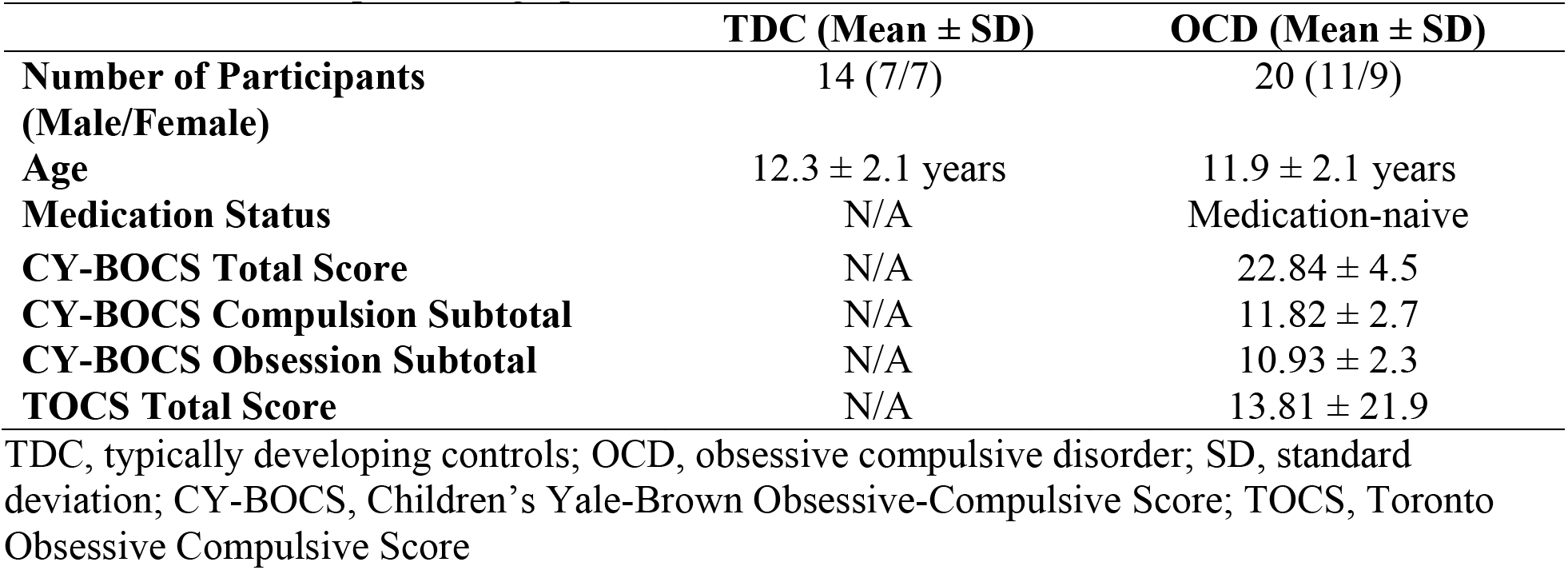
Overall Sample Demographic Characteristics

### 3.2 Go/No-Go task performance

There were no significant between-group differences in reaction time to the Go visual cue during the Go condition (t = <1.48, df = 28, p = 0.15), or to Go cues during the No-Go condition (t = −1.57, df = 28, p = 0.13). There were no significant between-group differences in the number of stop errors following the No-Go visual cue (t = −0.25, df = 28, p = 0.80) (see Table 2 for details).

**Table 2.** Behavioural Performance Across Reaction Time

|  | **TDC ms (±SD)** | **OCD ms (±SD)** | ***p*** | **Cohen's *d*** |
| --- | --- | --- | --- | --- |
| **Go: Mean Reaction Time** | 321 (± 62.9) | 390 (± 158.4) | 0.15 | 0.57 |
| **No-Go: Mean Reaction Time** | 423 (± 81.3) | 485 (± 122.5) | 0.13 | 0.59 |
| **Number of Errors** | 10.69 (± 10.68) | 11.76 (± 12.27) | 0.80 | 0.09 |

Mean reaction time, in milliseconds (ms ± SD), of button-press in response to Go visual cue in Go condition and incorrect button-presses in response to stop visual cue in No-Go condition. Number of errors in youth with OCD and TDC during No-Go trials. Cohen's *d* and *p* values represent effect size and significance of reaction times between-groups, respectively. TDC, typically developing controls; OCD, obsessive compulsive disorder; SD, standard deviation

### 3.3 Regional activation during task performance

Source Localization and Amplitude in Go Condition. Both groups showed activation in the primary visual cortex (V1) following presentation of the Go visual cue and the contralateral primary motor cortex (MI) following button-presses to the Go visual cue. There were no significant between-group differences in the latency or amplitude of V1 activity (t = 0.86, df = 28, p = 0.40) (Supplementary Figures 1 and 2). Average amplitude of MI activity at the time of the buttonpress was increased in the OCD versus TDC group (t = −2.35, df = 28, p = 0.03) (see Figure 1, Table 3).

**Figure 1.**
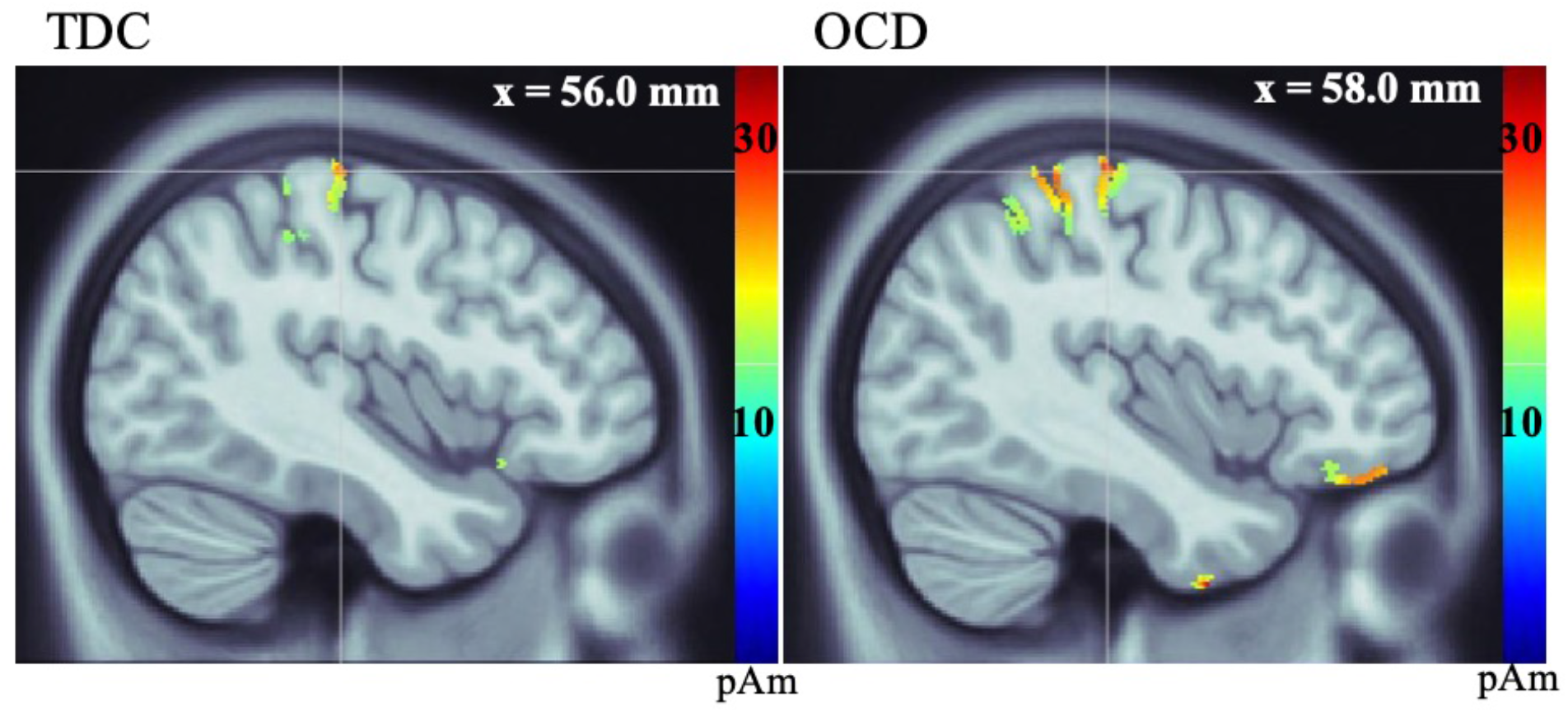
Source localization, measured in picoamperes (pAm), of group average response of primary motor cortex (MI) in TDC (left) and OCD (right) groups during button-press to Go visual cue.

**Table 3.** Latency and Amplitude of Activation During Go and No-Go Conditions

|  |  | Controls |  | OCD |  | *p* | Cohen's *d* |
| --- | --- | --- | --- | --- | --- | --- | --- |
| | | Latency<br>ms (±SD) | Amplitude<br>pAm (±SD) | Latency<br>ms (±SD) | Amplitude<br>pAm (±SD) | | |
| Go | V1 | 94.5<br>(± 11.1) | 64.3 (± 27.8) | 100.3<br>(± 18.5) | 55.9 (± 25.6) | 0.40 | 0.31 |
| | MI | N/A | 26.5 (± 14.2) | N/A | 46.9 (± 28.6) | 0.03* | 0.90 |
| No-Go | PCu | 164.0<br>(± 8.7) | 46.9 (± 22.2) | 97.4<br>(± 10.2) | 23.7 (± 8.7) | 0.001* | 1.38 |
| | OFC | 179.0<br>(± 9.1) | 31.7 (± 24.1) | 148.0<br>(± 6.43) | 21.3 (± 15.6) | 0.19 | 0.51 |
| | ACC | 178.0<br>(± 8.2) | 27.4 (± 32.0) | 141.7<br>(± 9.9) | 23.5 (± 10.4) | 0.72 | 0.16 |

Latency, measured in milliseconds (ms), and amplitude, measured in picoamperes (pAm), values for regions most active at the group level in the Go and No-Go conditions. Cohen's *d* and *p* values represent effect size and significance of amplitudes between-groups. OCD, obsessive compulsive disorder; V1, primary visual cortex; MI, primary motor cortex; PCu, precuneus; OFC, orbitofrontal cortex; ACC, anterior cingulate cortex; SD, standard deviation.
\* $p < 0.05$ for amplitude of activation

*Source Localization and Amplitude in No-Go Condition*. Both groups showed activation in V1 and precuneus (PCu) during successful stopping to the No-Go cue. No significant between-group differences in latency or amplitude of V1 activation were found. There was a significant between-group difference in amplitude of activation in the PCu, such that participants with OCD featured reduced amplitude at the group level, compared to TDC (t = 3.71, df =25, p = 0.001) (Supplementary Figures 3 and 4). In both groups, MI was active at the time of stop errors and the OFC was active after stop errors. There were no significant differences in MI. There was a trend towards reduced OFC amplitude in the OCD versus TDC group (t = 1.34, df = 25, p = 0.19). Average group activity of the ACC was only observed in the TDC group after stop errors. On further examination, ACC activity was found in 10 of 14 OCD participants (based on participant-level activations) following stop errors with no significant between-group differences found (t = 0.36, df = 21, p = 0.72) (see Figure 2, Table 3).

**Figure 2.**
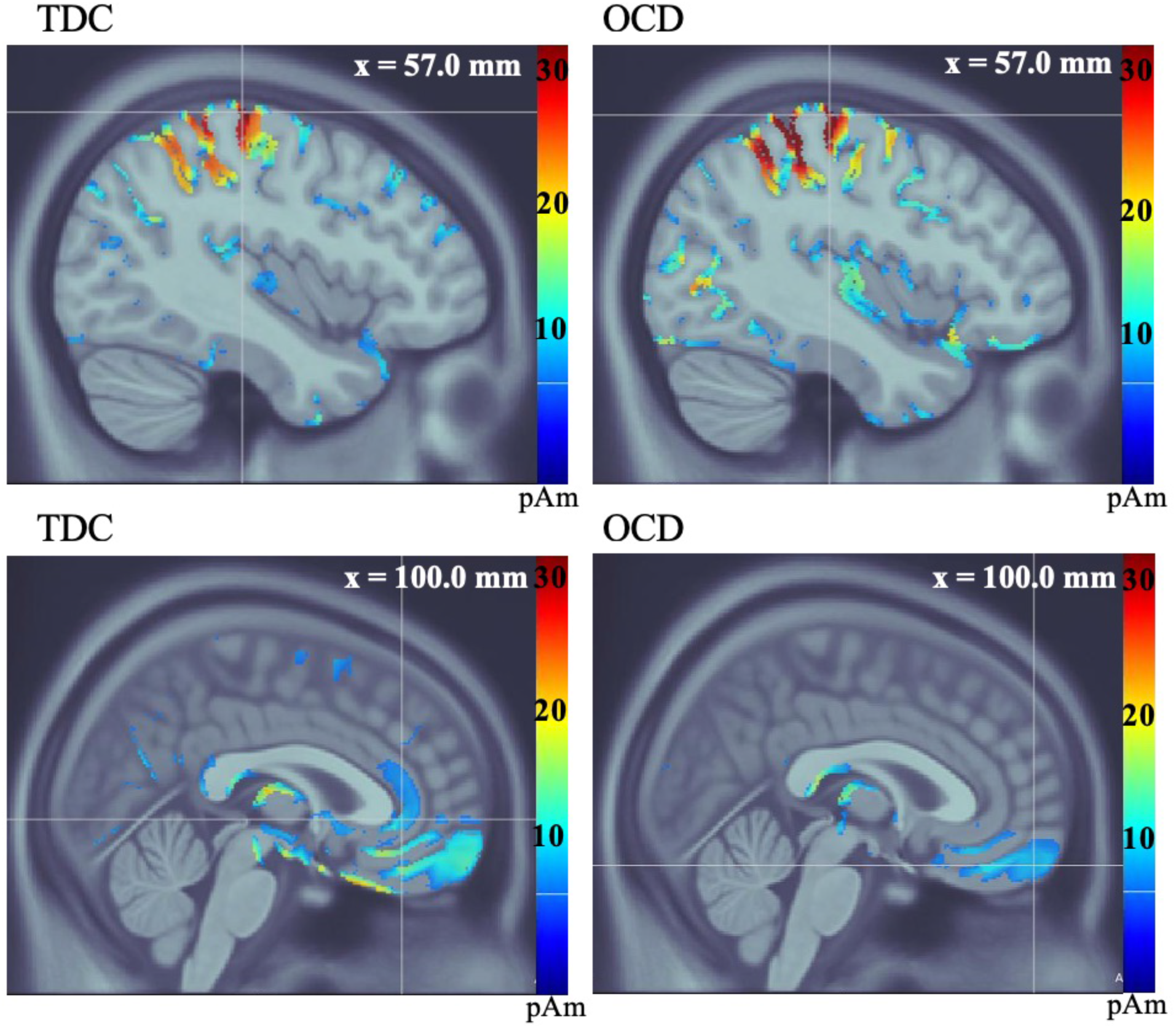
Average group source localization during incorrect button-press in response to stop visual cue in No-Go condition, measured in picoamperes (pAm). TDC group motor (MI) response (top left), and orbitofrontal and anterior cingulate cortices (bottom left). OCD group motor (MI) response (top right), and orbitofrontal cortex (bottom right). The OCD group does not show activation of the ACC at the group level (bottom right).

### 3.4 Post-hoc exploratory analyses

#### 3.4.1 Time-frequency responses within FST regions showing group differences (MI and OFC)

Group average TFR plots were generated and examined qualitatively by selecting for virtual sensors. MI activation during the Go condition and OFC activation during the No-Go condition were examined as these were areas of a priori interest that featured between-group differences. In the Go condition, the TFR plots for OCD in MI showed weak, ongoing theta oscillations throughout the trial and less ERS of alpha oscillations following button-press compared to TDC. In contrast, TFR plots for this region in the TDC group showed strong, ongoing theta oscillations for the entirety of the trial, and greater ERS of alpha oscillations following button-press (Figure 3). In the OFC for the No-Go condition, TFR plots for the OCD group showed continuous delta and theta oscillations throughout the trial and strong ERS in alpha oscillations following a stop error. In comparison, TFR plots for the TDC group showed consistent delta oscillation throughout the trial as well as transient ERS of beta oscillations and ERD of alpha oscillations following a stop error (Figure 4).

**Figure 3.**
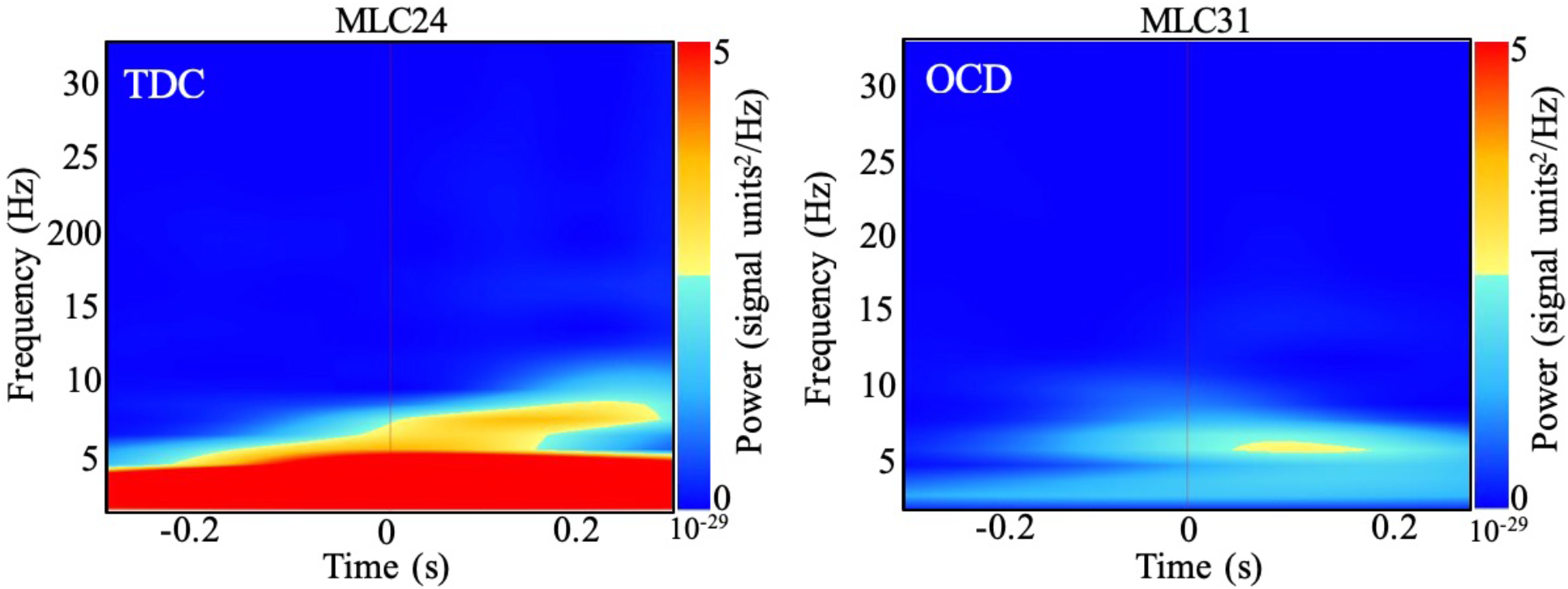
Average group time-frequency response of motor (MI) region activation following Go visual cue in TDC group (left) and OCD group (right).

**Figure 4.**
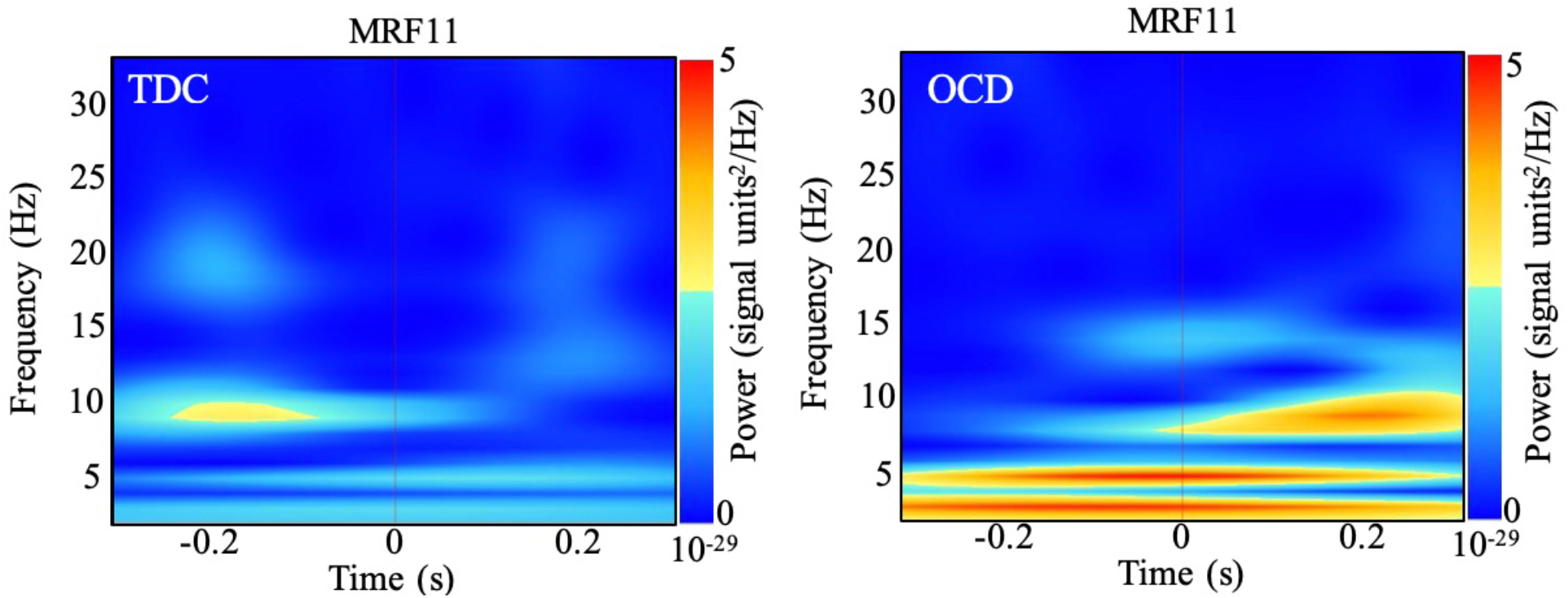
Average group time-frequency response of orbitofrontal cortex activation following button-press response to (A) No-Go visual cue in control group (B) and OCD group.

#### 3.4.2 Correlations between psychometric scores and amplitude of regional activation in MI and OFC

Exploratory correlation analysis was performed between amplitude of activation of regions of interest that differed between groups (i.e., MI and OFC). A trend towards a negative association between OFC and total CY-BOCS score was found (t = −1.86, df = 13, p = 0.09, r = −0.46). No significant correlation was found between amplitude of MI activation and total CY-BOCS scores (t = −0.08, df = 17, p = 0.94, r = −0.02).

## 4 Discussion

To our knowledge, this is one of the first studies to use MEG in children and youth with OCD and the only study measuring neural response during cognitive task performance in a medication-naïve sample. We did not find significant between-group differences on general Go/No-Go task performance. However, the total number of stop errors (following the No-Go visual cue) was numerically higher in the OCD group, consistent with prior findings of increased stop errors in youth with OCD during No-Go task performance (Bannon et al., 2002; Fitzgerald et al., 2010). In contrast, neural response during Go/No-Go task performance did differ significantly between OCD and TDC participants. During Go trials, we found increased MI activity corresponding to Go cue presentation and during No-Go trials, we found decreased PCu activity corresponding to No-Go cue presentation and a trend towards significantly reduced OFC amplitude during stop errors in OCD versus TDC. Our preliminary study in a small medication-naïve sample suggests that neural response within (and beyond) the FST circuit may be altered both during Go (attention/motor) and No-Go (response inhibition) task performance, and that MEG as an imaging tool may be sensitive to detecting such differences.

Higher amplitude of activation was observed in the MI region in OCD versus TDC at the time of Go cue presentation in the Go condition. Although not a core region of the FST, altered MI response in OCD is intriguing given the close connection between MI and the SMA, a key FST circuit region that has been implicated in OCD (de Wit et al., 2012). A recent meta-analysis of the fMRI literature found that both adults and youth with OCD showed hyperactivation of motor regions such as the SMA and pre-SMA, measured in tasks of response-inhibition, or task-switching (Norman et al., 2019). Increased excitability within MI, specifically, has also been shown in adults with OCD (Khedr et al., 2016), including in a study that found increased MI excitability during Go/No-Go Task performance in OCD that was also associated with an earlier (childhood) onset of OCD symptoms (Kang et al., 2019). A prior study suggested that increased activation in motor cortex in children and adults with OCD may be linked to increased difficulty in inhibiting responses and potentially related to compulsive behavior (Bhattacharjee et al., 2020). Importantly, MI has been a successful neural target for low-frequency (inhibitory) brain stimulation clinical trials aiming to treat OCD symptoms in adults that have not responded to medication or behavioural treatments (Rapinesi et al, 2019). Time-frequency analysis of MI following Go presentation revealed a qualitative pattern of decreased alpha band synchronization (i.e., a lesser increase in frequency of oscillations) following button-press to Go visual cue in OCD compared to TDC. A prior Go/No-Go MEG study found alpha oscillations increased in a non-clinical sample following either Go or No-Go cues, suggesting that increased alpha oscillations may signal successful attentional modulation to the cue to facilitate either a motor response or motor inhibition (Bauer et al., 2014; Hege et al, 2014). Increased MI amplitude and decreased alpha oscillatory activity may indicate suboptimal attentional modulation within this region during a simple visuomotor task (i.e., to Go cue) in medication-naïve children and youth with OCD.

Prior studies have implicated increased alterations within key frontal FST regions (i.e., activation of ACC and reduced activation of OFC) in OCD during response inhibition (Norman et al., 2019; Agam et al., 2014). In the current study, we found a trend towards reduced OFC amplitude during No-Go trials but no differences within the ACC. A prior MEG study also found altered OFC amplitude in participants with OCD, though that study showed increased (as opposed to decreased) OFC amplitude during a working memory task in adults with OCD compared to controls (Ciesielski et al., 2005). Differences in the direction of OFC alteration in the present (versus prior) MEG study may be due to differences between studies in participant age, task of interest, and factors related to medication exposure in the adult study. The time-frequency analysis revealed a pattern of increased OFC alpha band activity in OCD (as opposed to decreased in TDC), perhaps signalling increased attentional modulation in OCD following the No-Go cue (Hege et al., 2014). Therefore, increased alpha-related oscillatory activity may again signal suboptimal neural response during performance of a higher cognitive load task (i.e., No-Go). We also found a trend towards a negative association between OFC amplitude and total CY-BOCS score. A prior study found increased OFC activation in youth with OCD was associated with CYBOCS score improvement following treatment with SSRIs or cognitive behavioural therapy (Rubia et al., 2010). Preliminary findings from the current study and evidence from the prior literature suggests that OFC alterations may be present across pediatric and adult OCD during response inhibition, though the direction of findings may not be consistent, and may be influenced by age and/or medication exposure.

While both OCD and TDC groups engaged the PCu following No-Go cue presentation, the OCD group showed significantly lower amplitude of PCu activation compared to TDC. The PCu is a core region of the default-mode network (DMN) (Chen et al., 2019). Reduced network connectivity within the DMN has previously been found in adults with OCD. This lower amplitude of activation of the PCu in the OCD group found may potentially signal reduced efficiency or communication between the DMN and the executive control (task-positive) network which could impair cognitive tasks, such as task-switching and response inhibition, that typically evoke these networks (Chen et al., 2019).

A major strength of the current study is that investigation of a medication-naïve sample ensured that reported results are not driven or confounded by the effects of pharmacotherapeutic intervention on brain response (Bernstein et al., 2019). However, due to the challenge of recruiting medication-naïve participants at a tertiary care centre resulting in the small sample examined here, interpretation of our results must take into account the potential for inflated false positives in a small exploratory example. Other limits include the minimal characterization of the TDC sample and the wide age-range among study participants as MEG-measured neural response may be influenced by age-related differences in brain maturation affecting both processing speed and neural response (Ruhnau et al., 2011).

## 4.1 Conclusion

In conclusion, we present a novel investigation into the use of MEG to measure neural activity during response inhibition performance in a medication-naïve sample of children and youth with OCD. Our findings point to the potential for increased MI response during Go task performance and reduced PCu and OFC response during No-Go task performance. Future experiments should aim to recruit larger samples of children and youth with OCD to replicate findings, relate MEG-measured neural response to relevant behavioural and symptom domains and examine the effects of intervention.

## 6 Author contributions

EN completed all data processing and analysis, wrote the manuscript, and made the figures. CD assisted with data processing. JAEA assisted with statistical analyses. CD, ALW, TT, JAEA, SM, DJM, PDA, and SHA edited the manuscript.

