## Supplementary Figures for "Visuomotor Activation of Inhibition-Processing in Pediatric Obsessive Compulsive Disorder: A Magnetoencephalography Study"

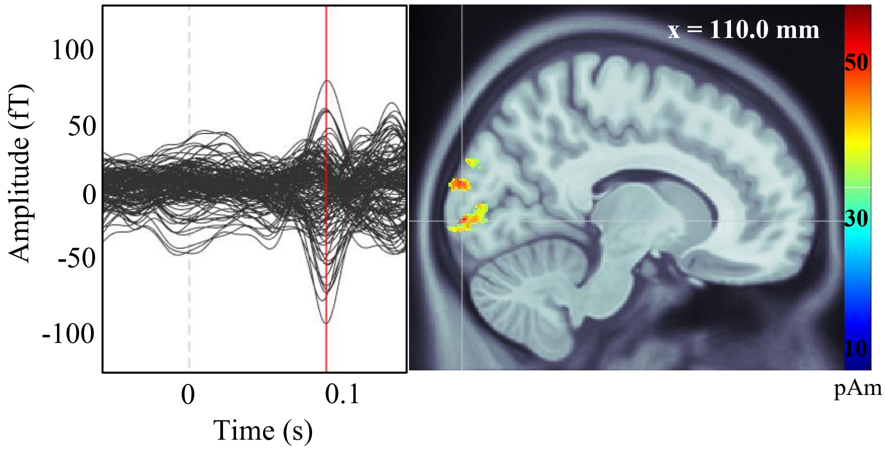


**Supplementary Figure 1.** Group average latency (left) measured in seconds, and source localization (right) measured in pAm, of visual (V1) response in the control group to visual cue in the Go condition.

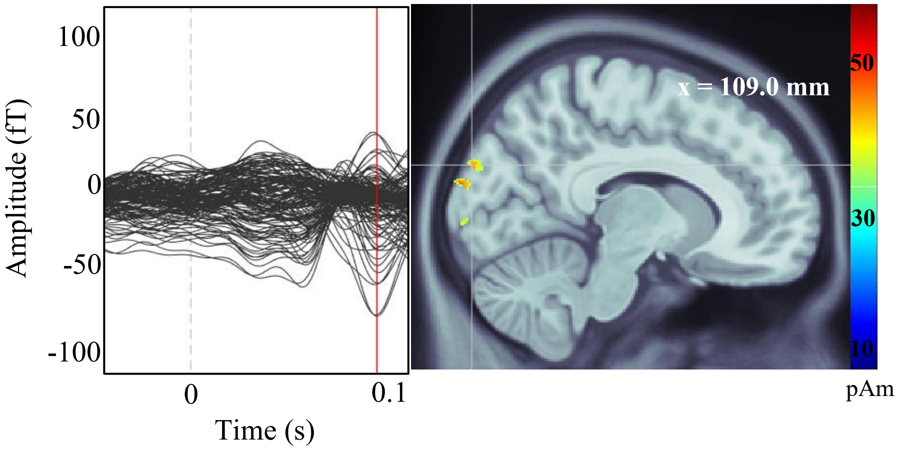


**Supplementary Figure 2.** Group average latency (left) measured in seconds (s), and source localization (right) measured in picoamperes (pAm), of visual (V1) response in the OCD group to visual cue in the Go condition.

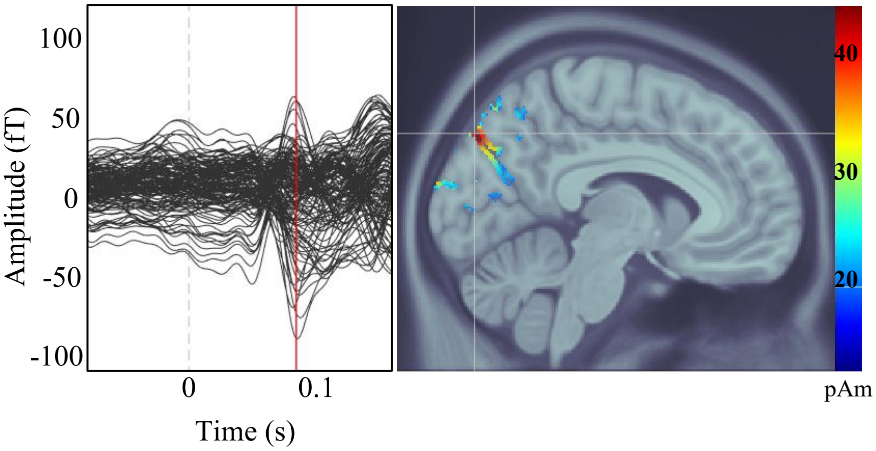


**Supplementary Figure 3.** Average group latency (left) measured in seconds (s), with red line depicting time of peak activation. Average group source localization (right), measured in picoamperes (pAm), of V1 and precuneus (middle) during visual response of control group in No-Go condition.

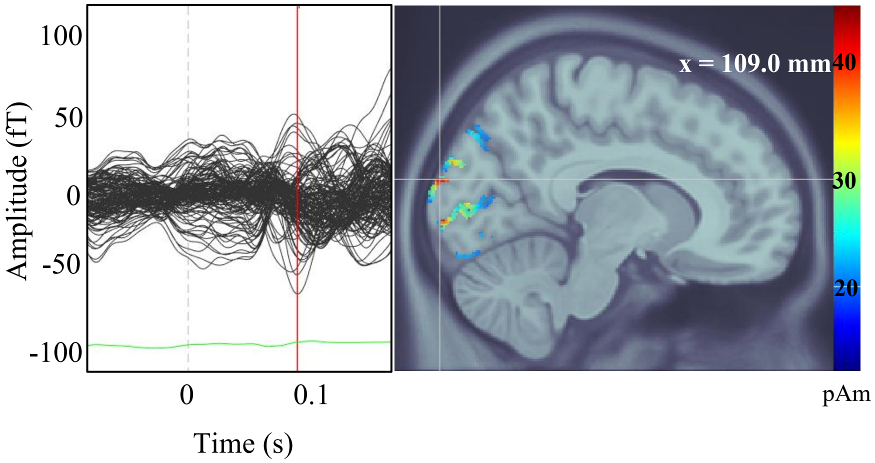


**Supplementary Figure 4.** Average group latency (left), measured in seconds (s), and average group source localization (right), measured in picoamperes (pAm), of V1 precuneus during visual response of OCD group to No-Go condition. The green line represents the global field power (GFP).
